# CancerSubminer: an integrated framework for cancer subtyping using supervised and unsupervised learning on DNA methylation profiles

**DOI:** 10.1101/2025.10.17.682936

**Authors:** Joung Min Choi, Liqing Zhang

**Affiliations:** Department of Computer Science Virginia Tech, Blacksburg, VA, US

## Abstract

Human cancer is highly heterogeneous, resulting in variable drug resistance and clinical outcomes. This complexity hinders accurate prognosis prediction and the development of targeted therapies. Molecular subtyping addresses these challenges by grouping cancers into more homogeneous subsets based on molecular characteristics, enabling subtype-specific treatment strategies. Subtyping is crucial for early diagnosis, personalized therapy, and improved survival by capturing differential therapeutic responses. Existing approaches to cancer subtyping fall into supervised and unsupervised categories. Supervised methods, often trained on The Cancer Genome Atlas (TCGA), rely on predefined subtype annotations but face limitations in generalizability and novel subtype discovery. Unsupervised methods, while capable of identifying new subtypes, may overlook widely recognized ones, hindering consistency with established classifications. Multi-omics approaches improve accuracy but are constrained by costs and data collection.

We propose CancerSubminer, a hybrid subtyping framework that integrates supervised and unsupervised learning. A subtype classifier is first trained on labeled data, after which clustering is applied to extracted features, with low-confidence samples reassigned to refine subtype boundaries. Model is retrained with the refined subtypes, and adversarial training corrects batch effects and learns domain-invariant features across labeled TCGA and unlabeled external datasets. A subsequent semi-supervised fine-tuning phase aligns subtypes between datasets and designates low-confidence samples as potential novel candidates. CancerSubminer was evaluated on five cancer types, including breast, bladder, brain, kidney, and thyroid cancers, using TCGA methylation data with annotated subtypes and unlabeled datasets from the Gene Expression Omnibus. The framework outperformed state-of-the-art subtyping models (iClusterPlus, iClusterBayes, NEMO) and clustering methods (Spectral, K-means). Kaplan–Meier survival analysis demonstrated significant prognostic separation (p < 0.05) for all cancers, including thyroid cancer where predefined subtypes showed no significance but CancerSubminer-derived subtypes did. These findings highlight CancerSubminer’s ability to identify distinct prognostic subtypes, mitigate batch effects, and improve prognostic stratification across heterogeneous datasets. CancerSubminer is publicly available at https://github.com/joungmin-choi/CancerSubminer.

## 1 Introduction

Cancer exhibits profound heterogeneity across multiple molecular and clinical dimensions, including genetic alterations, epigenetic modifications, transcriptomic programs, and phenotypic variability within and between patients [1]. This complexity drives diverse therapeutic responses and clinical outcomes, presenting significant challenges for accurate prognosis prediction and the development of effective targeted therapies. To address these challenges, extensive research efforts have focused on partitioning cancers into more homogeneous subgroups that better capture underlying biological mechanisms. This process, broadly termed cancer subtyping, has emerged as a cornerstone of precision oncology, enabling earlier diagnosis, personalized therapeutic approaches, and improved patient survival through subtype-specific treatment strategies.

Cancer subtype identification has evolved considerably from traditional morphological classification to molecular subtyping—an approach that groups tumors based on shared molecular characteristics rather than relying solely on histological features [2]. Genomic data have proven highly effective in subtyping studies across diverse cancer types. These molecular subtyping studies have successfully classified cancers into more uniform groups that demonstrate stronger correlations with clinical outcomes compared to conventional classification systems, thereby providing enhanced diagnostic, prognostic, and therapeutic strategies [3].

Early subtyping approaches were mostly supervised, relying on predefined labels and biomarker signatures to construct predictive classifiers. A representative example is PAM50 [4], which stratifies breast cancers into five intrinsic molecular subtypes using the expression profiles of 50 signature genes. Building on this foundation, machine learning methods extended supervised classification to incorporate genome-wide transcriptomic and epigenomic data. Although these strategies have shown good performance on datasets resembling their training cohorts, they also present certain limitations. They depend on predefined subtype annotations—often derived from The Cancer Genome Atlas (TCGA) [5]—and may therefore reflect biases present in those reference labels. In addition, supervised frameworks are limited in their capacity to uncover new subtypes that could be biologically or therapeutically relevant but are not represented in the training data.

In contrast, unsupervised methods such as clustering algorithms have been widely applied to enable de novo cancer subtype discovery. These approaches can uncover previously unknown molecular subgroups without reliance on predefined annotations, thereby facilitating novel subtype identification. However, unsupervised methods often fail to preserve well-established subtypes that carry clinical significance, leading to inconsistencies with existing diagnostic standards. Furthermore, clustering algorithms can be sensitive to noise and technical artifacts, compromising reproducibility across independent datasets.

To enhance robustness, multi-omics integration strategies have emerged that leverage complementary layers of biological information, including DNA methylation, gene expression, copy number variation, and proteomics data. Methods such as iClusterPlus [6], iClusterBayes [7], and NEMO [8] exemplify this approach, demonstrating improved accuracy and enabling deeper biological insights through comprehensive data integration. iClusterPlus extends the original iCluster framework by enabling integrative clustering of multi-omics data through a latent variable regression framework that jointly models diverse data types. By capturing shared latent oncogenic processes across data modalities, it facilitates more comprehensive cancer subtype discovery. iClusterBayes employs a fully Bayesian latent variable model that integrates omics data through data-type-specific likelihoods while incorporating Bayesian variable selection to identify driver features. Through MCMC-based inference, it provides posterior probabilities for cluster assignments. NEMO takes a network-based approach, developing a similarity algorithm that integrates patient-patient neighborhood graphs across omics layers and applies spectral clustering to the combined network. Despite these advances, the translation of multi-omics models into routine clinical practice remains constrained. Generating matched multi-omics datasets requires substantial computational resources, involves significant costs, and frequently suffers from missing data across modalities. Considering these limitations, the above studies have suggested that their frameworks could also be applied to single-omics data, although their performance in this setting has not been systematically assessed. These practical barriers highlight the continued need for subtyping approaches that can operate effectively with widely available single-omics platforms while maintaining potential clinical relevance.

DNA methylation has emerged as a particularly promising biomarker among single-omics platforms for cancer subtyping [9]. This key epigenetic modification regulates gene expression by adding methyl groups to CpG dinucleotides. Aberrant methylation patterns—including global hypomethylation and promoter hypermethylation of tumor suppressor genes—represent fundamental hallmarks of tumorigenesis. Distinct methylation landscapes effectively delineate cancer subtypes across diverse tumor types, often providing prognostic and therapeutic insights that complement or even surpass those derived from transcriptomic data. Nevertheless, integrating methylation data from multiple cohorts introduces significant challenges. Batch effects stemming from technical differences across platforms or laboratories can mask genuine biological variation and introduce bias into subtype discovery [10]. Therefore, when performing subtyping with multi-source data, it becomes critical to identify and correct for these batch effects to ensure that discovered subtypes reflect underlying biological variation and clinical relevance rather than technical artifacts. Batch correction is essential during cancer subtyping to enhance reproducibility across studies and improve the reliability.

To address these limitations, we propose CancerSubminer, a cancer subtyping framework that integrates supervised and unsupervised learning. The framework begins with a supervised phase, where a subtype classifier is trained on labeled datasets to capture widely recognized subtype categories. The learned feature representations are then clustered, and samples with low classification confidence are reassigned to refine subtype boundaries and to flag potential novel subtypes. The model is subsequently retrained with these refined labels, while adversarial training is applied to mitigate batch effects and enforce domain-invariant representations across both labeled and unlabeled datasets. A semi-supervised fine-tuning step further aligns subtypes between datasets and facilitates the discovery of additional candidate subgroups. CancerSubminer was evaluated across five cancer types—breast, bladder, brain, kidney, and thyroid—using large-scale DNA methylation datasets. Benchmarking against state-of-the-art subtyping frameworks (iClusterPlus, iClusterBayes, NEMO) and clustering methods (spectral clustering, K-means) demonstrated superior performance in terms of clustering quality, batch effect correction, and subtype alignment across datasets. Kaplan–Meier survival analyses revealed significant prognostic separation in all five cancer types, including thyroid cancer, where predefined subtypes failed to show prognostic relevance. Clinical enrichment analyses further supported the clinical validity of the identified subtypes. Collectively, these results establish CancerSubminer as a robust and versatile framework for cancer subtyping, capable of refining established classifications, correcting for batch effects, and improving prognostic stratification across heterogeneous cancer cohorts.

## 2 Methods

Let *n*_*s*_ and *n*_*t*_ denote the numbers of samples in the source and target datasets, respectively, and let *m* represent the number of CpG sites shared across all data. The source DNA methylation data matrix is given by 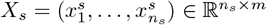, where each sample is annotated with established subtype labels. The target methylation dataset, defined as 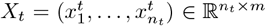, consists of multiple unlabeled cohorts. We assume *X*_*s*_ and *X*_*t*_ arise from distinct but related distributions, reflecting heterogeneity across cohorts due to technical batch effects and biological variability.

The proposed CancerSubminer framework is organized into four phases: (1) supervised pre-training of a subtype classifier, (2) clustering with confidence-based reassignment to refine subtype boundaries, (3) adversarial training to achieve domain-invariant feature representations, and (4) semi-supervised fine-tuning with subtype alignment to harmonize source and target distributions (Fig. 1).

**Figure 1:**
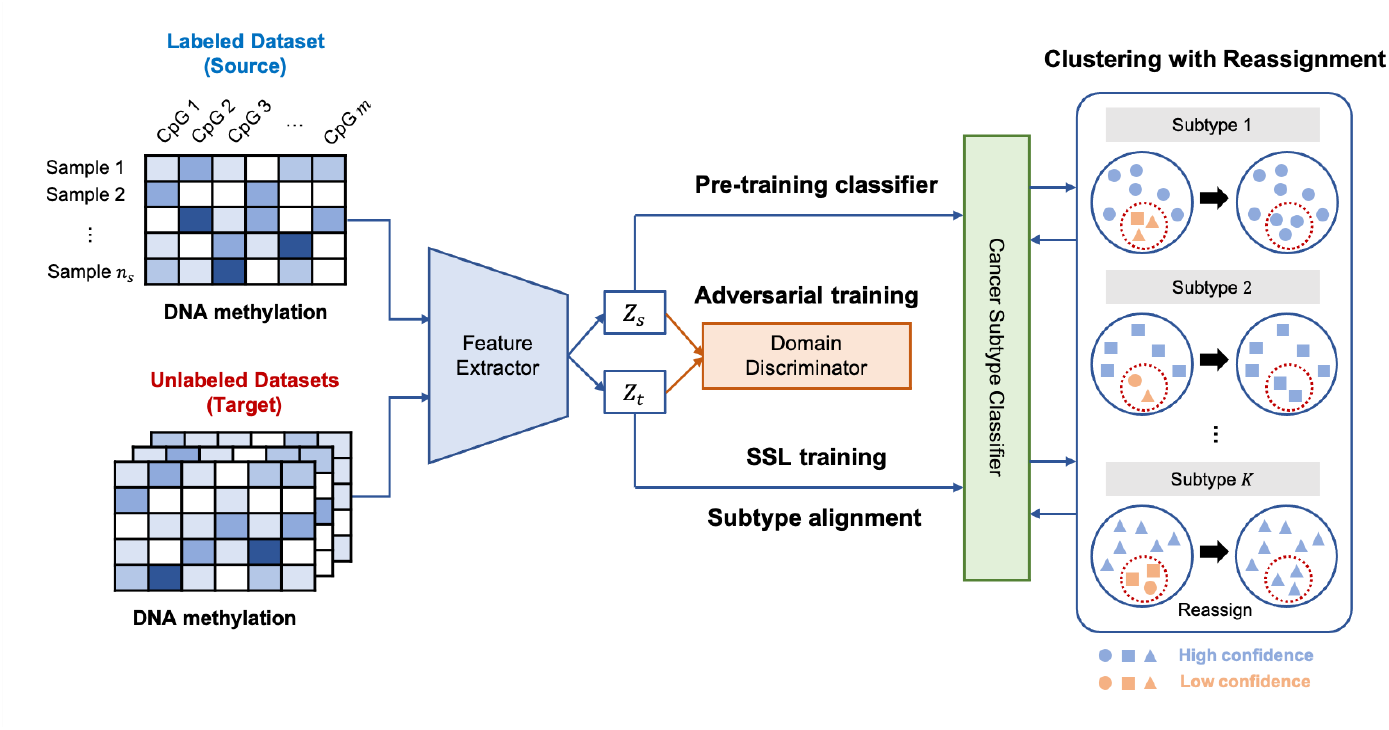
Overall workflow of the CancerSubminer framework. The method proceeds through four sequential phases: supervised pre-training, clustering with reassignment, adversarial domain adaptation, and semi-supervised fine-tuning with subtype alignment.

### 2.1 Supervised pre-training of the subtype classifier

The first stage aims to initialize the model by learning discriminative representations of predefined subtypes. A feature extractor *F*_*θ*_ and subtype classifier *C*_*ϕ*_ are trained jointly on the labeled source dataset *X*_*s*_. The feature extractor consists of fully connected layers with batch normalization and nonlinear activations, while the classifier module ends with a softmax layer to produce subtype posterior probabilities. The objective is to minimize the subtype classification error through the cross-entropy loss:

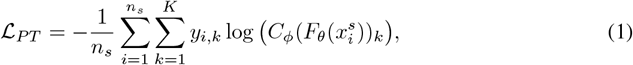

where *K* is the number of predefined subtypes, *y*_*i*,*k*_ is a binary indicator of the ground-truth subtype, and 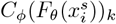 represents the predicted probability that sample *i* belongs to subtype *k*. This supervised pre-training ensures that the learned features are aligned with widely accepted subtype definitions, forming a baseline representation before integration with unlabeled data.

### 2.2 Clustering with confidence-based reassignment

While pre-training captures known subtypes, it does not fully account for potential heterogeneity within or across cohorts. To refine subtype definitions and identify candidate novel groups, unsupervised clustering was applied to the embeddings produced by *F*_*θ*_. K-means clustering is employed due to its efficiency, where the number of clusters *K* is either provided by the user or automatically estimated by progressively increasing from two fewer than the number of predefined subtypes until the silhouette score no longer improves.This adaptive search facilitates the discovery of finer-grained subtype structures that extend beyond the predefined subtype definitions.

To address inconsistencies between classifier predictions and cluster memberships, we incorporate a confidence-based reassignment strategy. Each sample’s posterior probability vector is obtained from *C*_*ϕ*_. Samples with maximum predicted probability below a confidence threshold (*τ* = 0.95) are considered ambiguous and reassigned to the majority subtype among high-confidence samples in the same cluster. This process ensures stability by relying on reliable predictions to guide reassignment. After reassignment, the model is retrained with the updated labels, and the clustering–reassignment cycle is repeated until assignments converge. This iterative refinement simultaneously sharpens subtype boundaries and highlights samples that may represent emerging subgroups not captured by the initial classification.

### 2.3 Adversarial training for domain-invariant feature learning

Differences between *X*_*s*_ and *X*_*t*_ arise from both biological diversity and systematic batch effects, which can hinder the transferability of subtype classifiers. To overcome this, CancerSubminer employs adversarial training to learn domain-invariant features. A domain discriminator *D*_*ψ*_ is introduced to distinguish whether embeddings are derived from the source or target domain. It consists of multiple dense layers with softmax output. During training, *D*_*ψ*_ is optimized to correctly classify the domain origin, while *F*_*θ*_ is simultaneously optimized to produce embeddings that confuse *D*_*ψ*_. This adversarial process encourages the distributions of *X*_*s*_ and *X*_*t*_ to align, thereby mitigating batch effects. The adversarial domain loss is defined as:

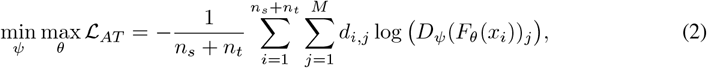

where *M* is the number of domains, *d*_*i*,*j*_ denotes the domain label, and *D*_*ψ*_(*F*_*θ*_(*x*_*i*_))_*j*_ is the predicted probability of sample *i* belonging to domain *j*. By iteratively updating *ψ* and *θ*, the model achieves a balance where *F*_*θ*_ extracts features that are discriminative for subtype classification yet invariant to dataset origin.

### 2.4 Semi-supervised fine-tuning with subtype alignment

The final phase of CancerSubminer integrates semi-supervised learning (SSL) with subtype alignment to enhance model generalization and ensure consistent subtype definitions across domains. This step both stabilizes pseudo-label assignments and identify potential novel subtypes in target cohorts.

SSL begins by assigning pseudo-labels *ŷ_j_* to each target sample 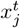 according to the subtype with the highest posterior probability predicted by the classifier. These pseudo-labels are updated iteratively during training. The feature extractor and subtype classifier are optimized using a weighted crossentropy loss applied to both the labeled source data and the pseudo-labeled target data:

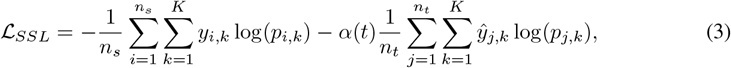

where *p*_*i*,*k*_ and *p*_*j*,*k*_ denote the predicted subtype probabilities for source and target samples, respectively. The coefficient *α*(*t*) increases gradually with epochs, balancing the influence of target pseudo-labels to prevent overfitting to noisy assignments in early training [11, 12].

However, pseudo-label assignments are inherently prone to errors, particularly when target samples lie near decision boundaries. Incorrect pseudo-labels can propagate noise, causing the classifier to reinforce spurious patterns and weakening generalization. To address this, CancerSubminer incorporates clustering with reassignment in every SSL epoch. Specifically, embeddings of both source and target samples are clustered using the *K* determined in the second phase. Target samples whose pseudo-labels conflict with the dominant subtype of their cluster are reassigned to the majority subtype of source samples in that cluster. This mechanism mitigates label noise by leveraging the stability of source-labeled samples as anchors for cluster consistency.

After multiple SSL iterations have stabilized pseudo-label assignments, the model applies subtype alignment to explicitly reduce discrepancies across domains. The goal is to minimize batch distinctions by ensuring that embeddings corresponding to the same subtype form compact clusters across both source and target domains. For each subtype *k*, centroids are computed separately for the source and target:

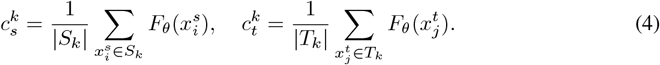

The subtype alignment loss is then defined as:

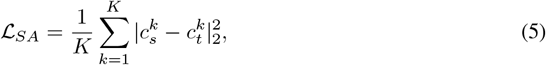

which encourages embeddings from the same subtype to converge while aligning subtypes across domains.

A key distinction from previous supervised subtyping framework is that CancerSubminer allows for the emergence of novel subtype candidates. Since target datasets may contain subgroups absent from the source data, alignment is restricted to high-confidence samples, preventing unstable assignments from distorting centroids. Low-confidence samples are grouped separately and monitored for signs of forming consistent clusters. If adding such a cluster improves both the silhouette score and classification performance, it is designated as a candidate novel subtype. Through this iterative process, fine-tuning simultaneously consolidates known subtype definitions, harmonizes embeddings across domains, and remains sensitive to the discovery of new subgroup structures in heterogeneous cancer cohorts.

#### 3 Experimental Design

### 3.1 Data collection

We obtained source datasets from The Cancer Genome Atlas (TCGA) [5], which provides DNA methylation profiles from solid primary tumor samples generated using Illumina Human Infinium 450K and 27K arrays. Our study focused on five cancer types: breast invasive carcinoma (BRCA), bladder urothelial carcinoma (BLCA), glioblastoma multiforme (GBM), thyroid carcinoma (THCA), and kidney renal clear cell carcinoma (KIRC). We obtained reference subtype annotations for each cancer type from previously established studies [13, 14, 15, 16, 17]. To build unlabeled target cohorts, we additionally retrieved three independent DNA methylation datasets per cancer type from the Gene Expression Omnibus (GEO) [18]. These GEO datasets did not include subtype labels and were therefore incorporated as unlabeled domains in our evaluation. An overview of dataset composition, including sample sizes for each cohort is provided in Table 1.

**Table 1:** Datasets used for BCtypeFinder evaluation.

| Cancer | # of subtypes | Source Dataset | # of samples | Target Dataset | # of samples |
| --- | --- | --- | --- | --- | --- |
| BRCA | 5 | TCGA-BRCA | 1060 | GSE69914, GSE75067, GSE72245 | 611 |
| BLCA | 4 | TCGA-BLCA | 418 | GSE183920, GSE193208, GSE50409 | 959 |
| GBM | 5 | TCGA-GBM | 375 | GSE200647, GSE195640, GSE195684 | 472 |
| THCA | 4 | TCGA-THCA | 496 | GSE97466, GSE72729, GSE86961 | 156 |
| KIRC | 5 | TCGA-KIRC | 441 | GSE113501, GSE61441, GSE105260 | 222 |

### 3.2 Data preprocessing

Data preprocessing was performed in line with established approaches [19, 20]. To ensure comparability across cohorts, only CpG sites shared between the source and target datasets were retained. CpG sites with more than 20% missing values were excluded to minimize bias during training, and the remaining missing values were imputed using the median. For dimensionality reduction, CpGs were grouped into 3,000 clusters using K-means clustering, and the representative feature for each cluster was defined as the median beta value.

## 4 Results

### 4.1 Assessing subtype compactness and separation across methods

We first assessed the subtyping performance of CancerSubminer by evaluating both intra-subtype compactness and inter-subtype separation. Silhouette score and Davies–Bouldin index (DBI) were calculated on the source datasets across five cancer types and compared against multiple subtyping strategies. Specifically, CancerSubminer was benchmarked against three state-of-the-art subtyping frameworks—iClusterPlus, iClusterBayes, and NEMO—which were originally proposed for multiomics integration but have also been extended by their authors for application to single-omics data. Widely used machine learning–based clustering approaches (spectral clustering and K-means) were also included as baselines. The optimal number of subtypes for each approach was determined either by the procedures recommended by the respective authors or, in the case of machine learning methods, by selecting the solution with the highest Silhouette score (Supplementary Material S1). In addition, clustering quality was compared against established subtype annotations available for the source datasets (denoted as ‘original’).

As shown in Table 2, CancerSubminer consistently outperformed the predefined subtypes, achieving higher silhouette scores and lower DBI across all cancer types. For BRCA, BLCA, and KIRC, competing methods also demonstrated improvements over established subtypes; however, CancerSubminer delivered the most robust performance overall, with the exception of DBI in BLCA. To further interpret these results, we applied Uniform Manifold Approximation and Projection (UMAP) [21]. For CancerSubminer, UMAP was performed using the extracted features, whereas for the comparison methods, the preprocessed input data were projected into two dimensions and labeled by batch and predicted subtype. For established subtypes, visualization was limited to datasets with available annotations. As illustrated in Fig. 2, Fig. 3, and Supplementary Material S2, CancerSubminer produced well-separated and coherent clusters, while mitigating batch-driven variation. In BLCA, NEMO generated numerous small clusters, indicative of potential over-clustering, which likely improved DBI through increased separation but reduced the silhouette score by misplacing samples near inappropriate cluster boundaries (Fig. 2). By contrast, CancerSubminer maintained balanced separation and compactness. Moreover, iClusterPlus and NEMO showed residual batch effects, with subtypes segregating within individual batches rather than spanning across them, particularly in BLCA and THCA. Interestingly, original subtype labels exhibited weak separation in most cancers except BRCA. iClusterBayes eliminated overt batch structure but failed to yield distinct subtype separation. Collectively, these results highlight CancerSubminer’s ability to refine subtype boundaries, correct batch-specific biases, and align subtype distributions across heterogeneous cohorts.

**Table 2:** Comparison of subtyping performance between the proposed and baseline methods using clustering metrics for the TCGA (source) dataset.

| Cancer |  | Original | CancerSubminer | iClusterPlus | iClusterBayes | NEMO | Spectral | K-means |
| --- | --- | --- | --- | --- | --- | --- | --- | --- |
| BRCA | # of subtypes | 5 | 5 | 3 | 6 | 11 | 4 | 3 |
|  | Silhouette score | 0.242 | **0.464** | 0.271 | -0.034 | 0.177 | -0.007 | 0.276 |
|  | DBI | 1.377 | **0.739** | 1.382 | 39.7 | 1.278 | 15.649 | 1.365 |
| BLCA | # of subtypes | 5 | 5 | 6 | 4 | 10 | 4 | 4 |
|  | Silhouette score | 0.176 | **0.421** | 0.121 | -0.051 | 0.296 | 0.078 | 0.388 |
|  | DBI | 2.377 | 0.904 | 1.710 | 18.258 | **0.787** | 3.478 | 1.366 |
| GBM | # of subtypes | 5 | 4 | 3 | 4 | 4 | 3 | 4 |
|  | Silhouette score | 0.553 | **0.628** | 0.194 | -0.046 | 0.379 | -0.013 | 0.392 |
|  | DBI | 1.405 | **0.508** | 1.322 | 34.81 | 0.769 | 5.295 | 0.753 |
| THCA | # of subtypes | 5 | 3 | 3 | 6 | 5 | 3 | 3 |
|  | Silhouette score | 0.591 | **0.685** | 0.341 | -0.066 | 0.259 | 0 | 0.369 |
|  | DBI | 0.586 | **0.481** | 1.253 | 18.538 | 1.173 | 6.52 | 1.055 |
| KIRC | # of subtypes | 4 | 2 | 2 | 5 | 10 | 4 | 3 |
|  | Silhouette score | -0.056 | **0.704** | 0.079 | -0.084 | 0.256 | 0.087 | 0.558 |
|  | DBI | 5.914 | **0.466** | 2.98 | 49.029 | 1.517 | 1.669 | 0.858 |

**Figure 2:**
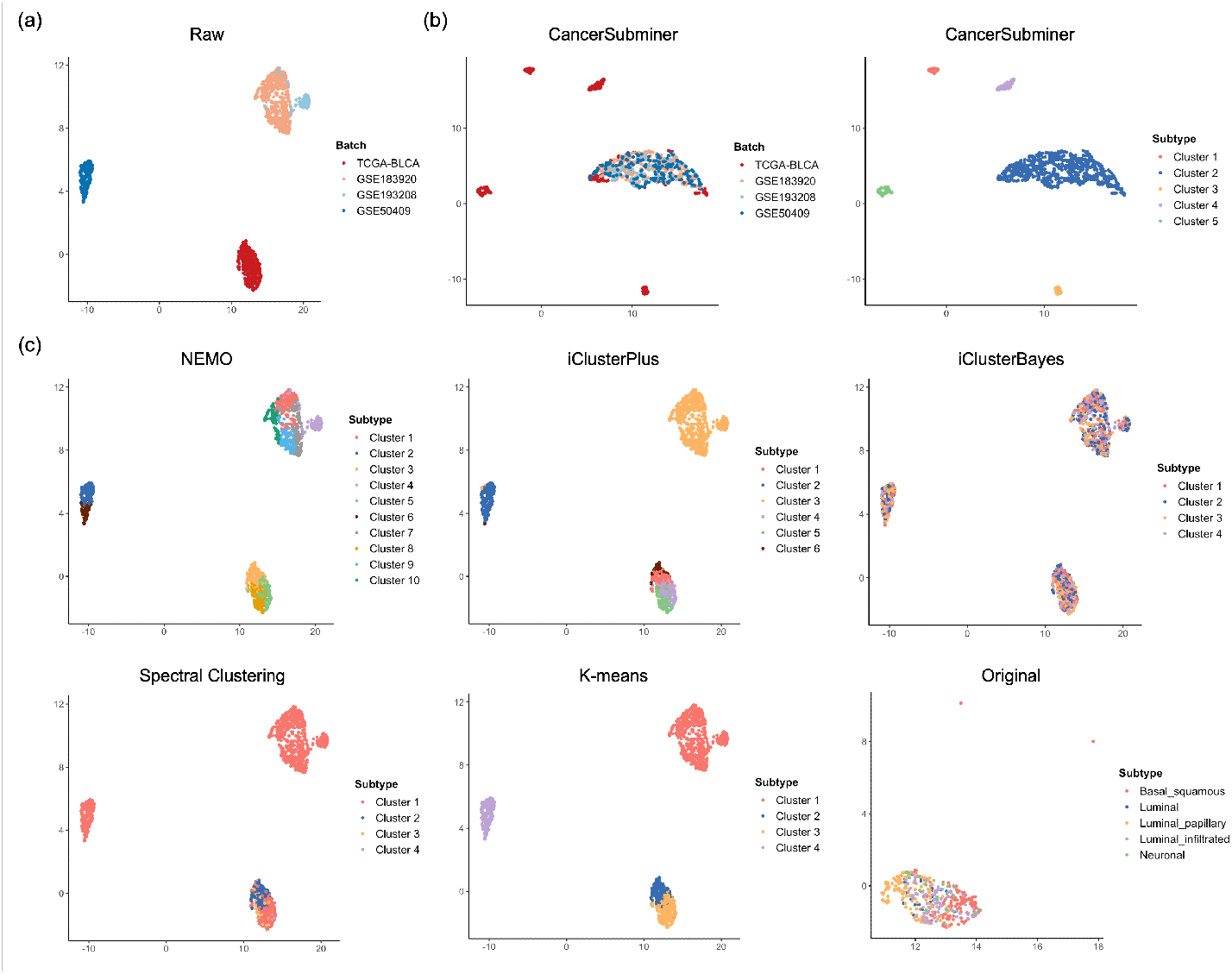
UMAP visualization of BLCA source and target datasets. (a) UMAP plot of the uncorrected dataset, colored by batch. (b) UMAP plots of features extracted by CancerSubminer, colored by batch (left) and by identified cancer subtypes (right). (c) UMAP plots of subtype assignments from comparison methods, including NEMO, iClusterPlus, iClusterBayes, Spectral Clustering, K-means, and the original dataset annotations.

**Figure 3:**
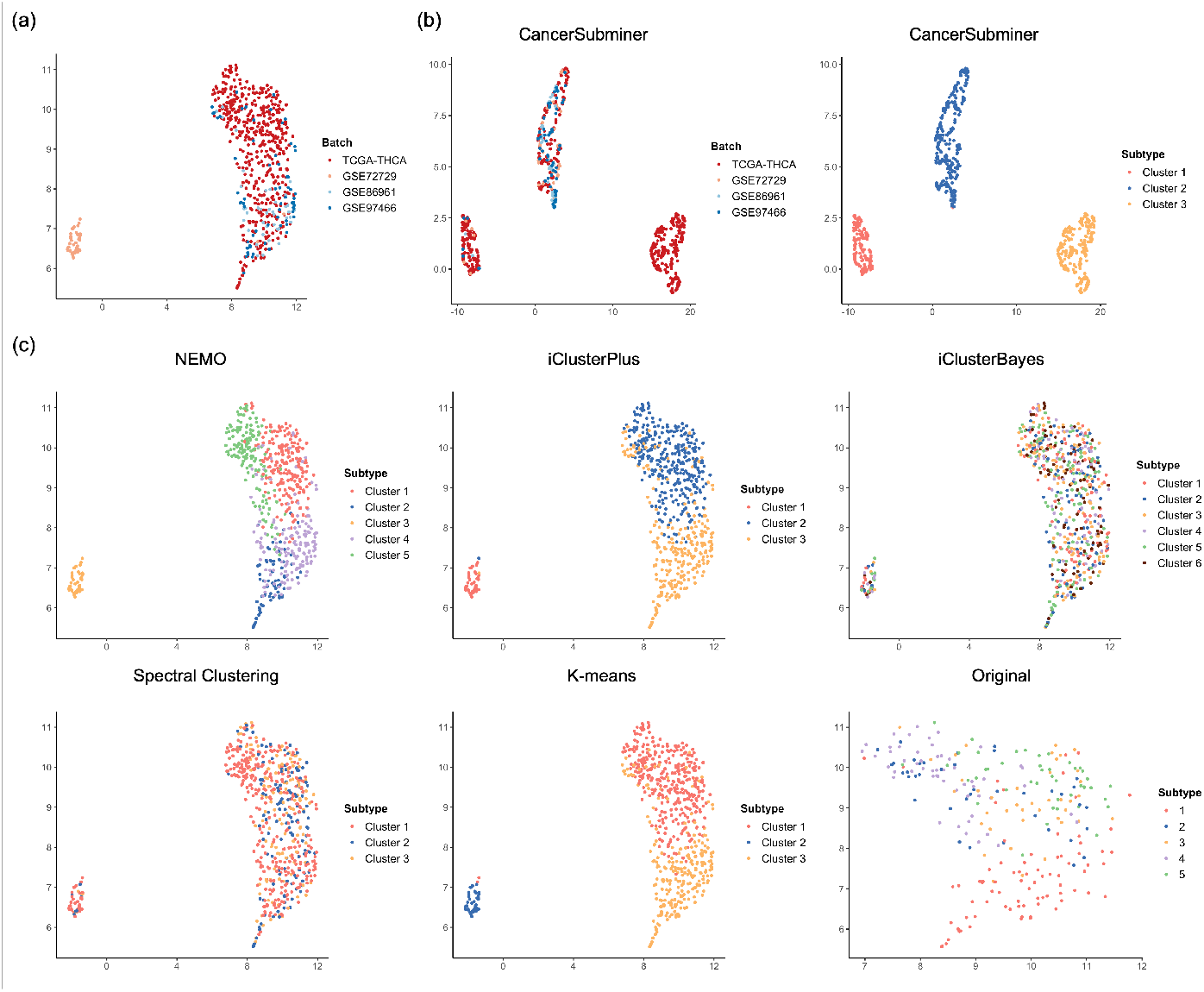
UMAP visualization of the source and target datasets in THCA. (a) UMAP representation of the uncorrected dataset, colored by batch. (b) UMAP plots of features extracted by CancerSubminer, colored by batch (left) and by identified cancer subtypes (right). (c) UMAP plots illustrating subtype assignments generated by alternative methods, including NEMO, iClusterPlus, iClusterBayes, Spectral Clustering, and K-means, alongside the original dataset annotations.

### 4.2 Investigating the prognostic significance of identified subtypes

A primary objective of cancer subtyping is to stratify patients into coherent groups that share similar prognoses yet exhibit distinct clinical outcomes across subtypes. Effective subtyping enhances the ability to predict survival probability and treatment response, as patients in different subtypes often benefit from distinct therapeutic strategies. To assess the prognostic relevance of the identified subtypes, we performed Kaplan–Meier (KM) survival analysis with log-rank testing using source datasets that contain overall survival information. For BRCA, we additionally evaluated one target cohort (GSE75067), which also includes survival data.

The results (Fig. 4, Fig. 5, Supplementary Material S3, S4) show that subtypes identified by CancerSubminer consistently achieved statistically significant survival separation across all five cancer types (log-rank test, *p <* 0.05). In BRCA, both the established subtypes and those identified by CancerSubminer demonstrated significant prognostic differences in TCGA ((Fig. 4) and GSE75067 (Supplementary Material S3), whereas most comparison methods failed to achieve significance.

**Figure 4:**
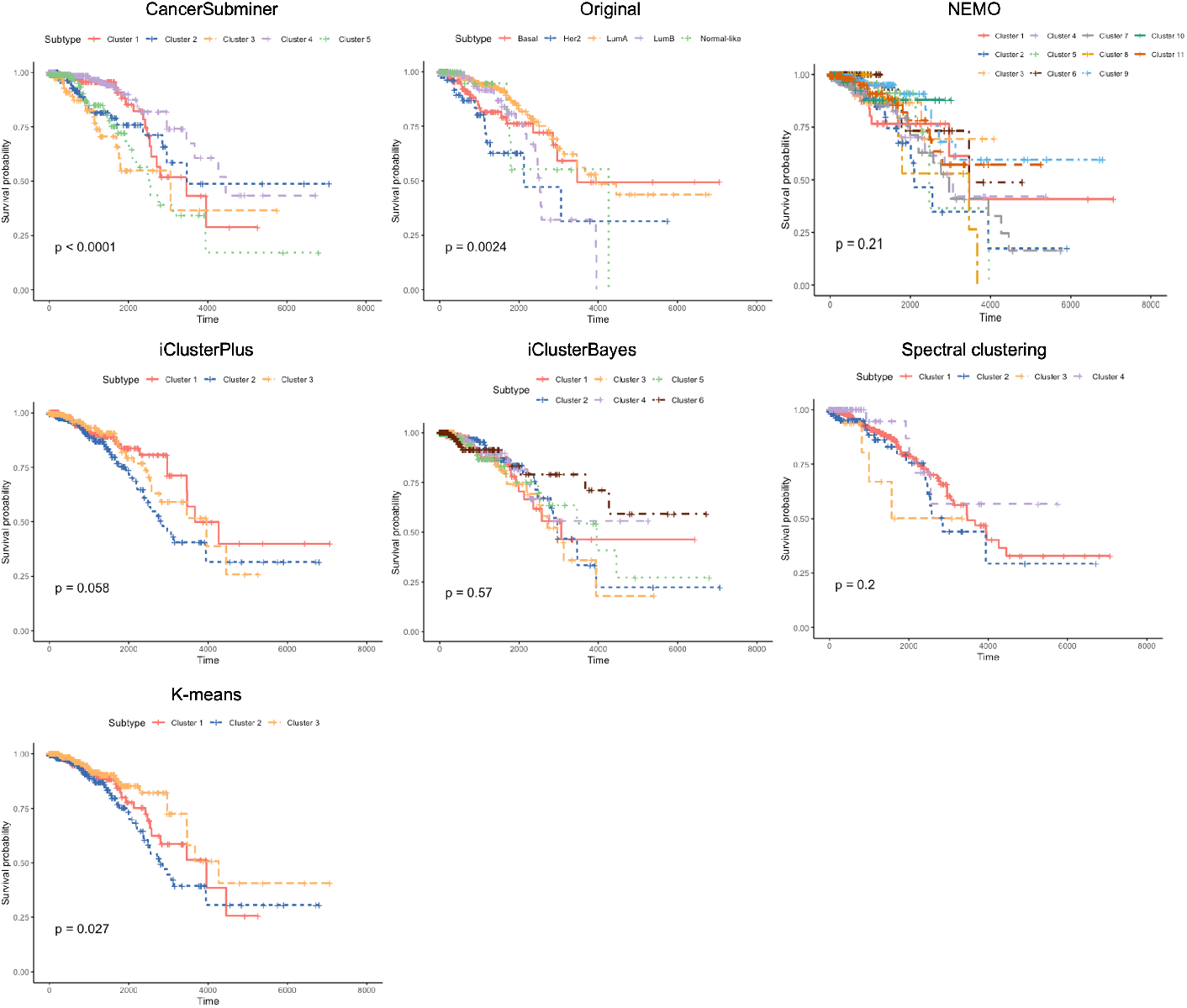
Kaplan–Meier survival analysis of BRCA subtypes identified from the TCGA dataset, with log-rank test results comparing CancerSubminer against baseline clustering methods.

**Figure 5:**
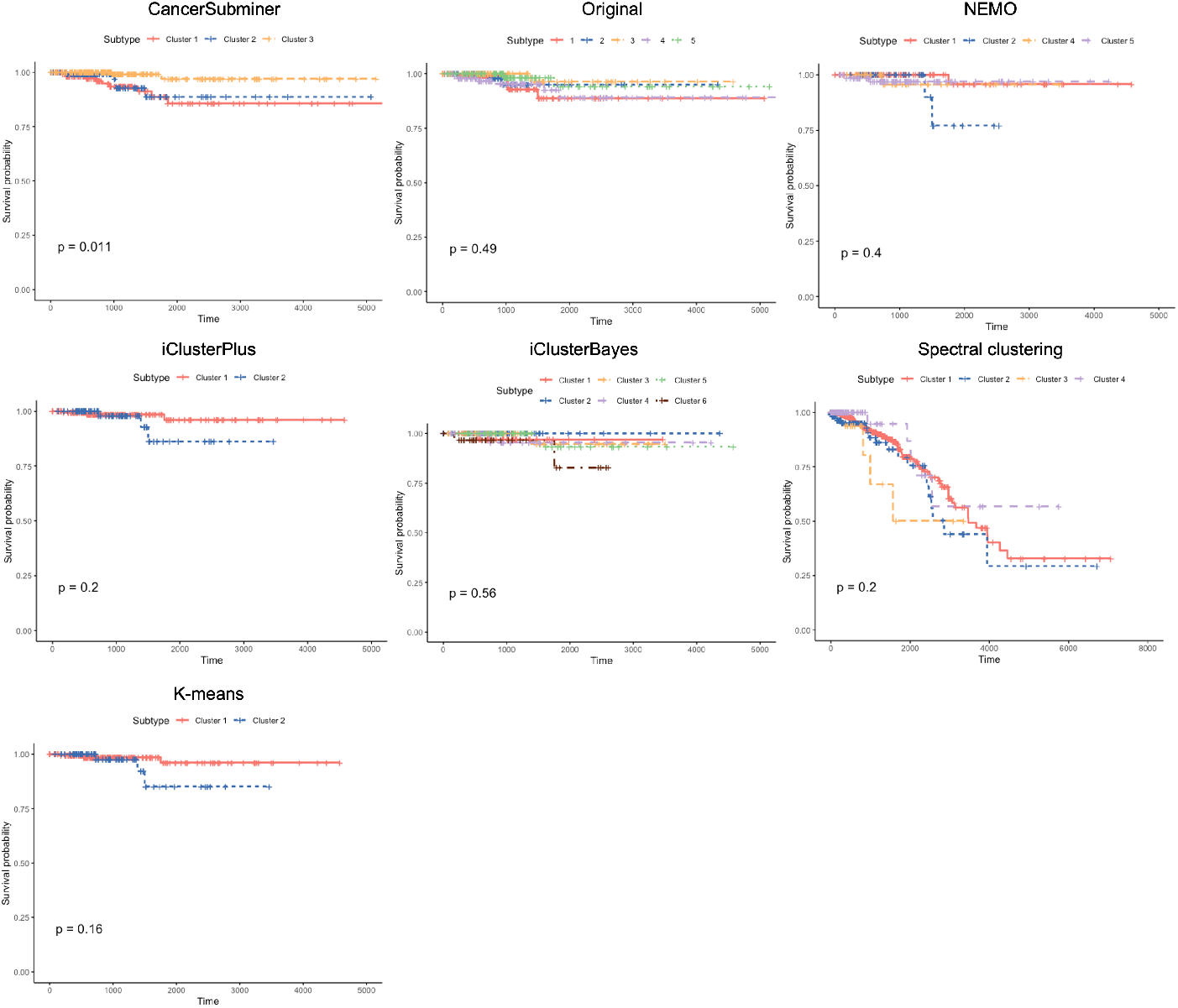
Kaplan–Meier survival analysis of THCA subtypes identified from the TCGA dataset with log-rank test results.

Among the alternative frameworks, NEMO performed best, yielding significant separation in BLCA, GBM, and KIRC (Supplementary Material S4). Notably, in THCA, established subtypes showed no significant prognostic distinction, yet the subtypes defined by CancerSubminer exhibited clear and statistically significant differences (Fig. 5). These findings highlight the ability of CancerSubminer to capture clinically meaningful heterogeneity beyond existing subtype definitions, underscoring its potential to improve prognostic stratification and inform treatment decision-making.

### 4.3 Evaluation of clinical associations across subtypes

To evaluate the clinical relevance of the subtyping results, we analyzed clinical variables that characterize differences between subtypes within each cancer type and performed enrichment testing. To ensure consistency and avoid bias, we selected the same set of clinical parameters across all five cancers: age at initial diagnosis and four pathological variables describing tumor progression (pathologic T), lymph node involvement (pathologic N), metastasis (pathologic M), and overall stage (pathologic stage). For each cancer type, enrichment was tested on the clinical parameters available in its dataset. Statistical significance was assessed using the chi-squared test for independence for categorical variables and the Kruskal–Wallis test for continuous variables. In the results (Table 3), “ns” indicates non-significance, while *, **, and *** denote *p <* 0.05, *p <* 0.01, and *p <* 0.001, respectively. CancerSubminer consistently delivered the strongest associations across all five cancers.

**Table 3:**
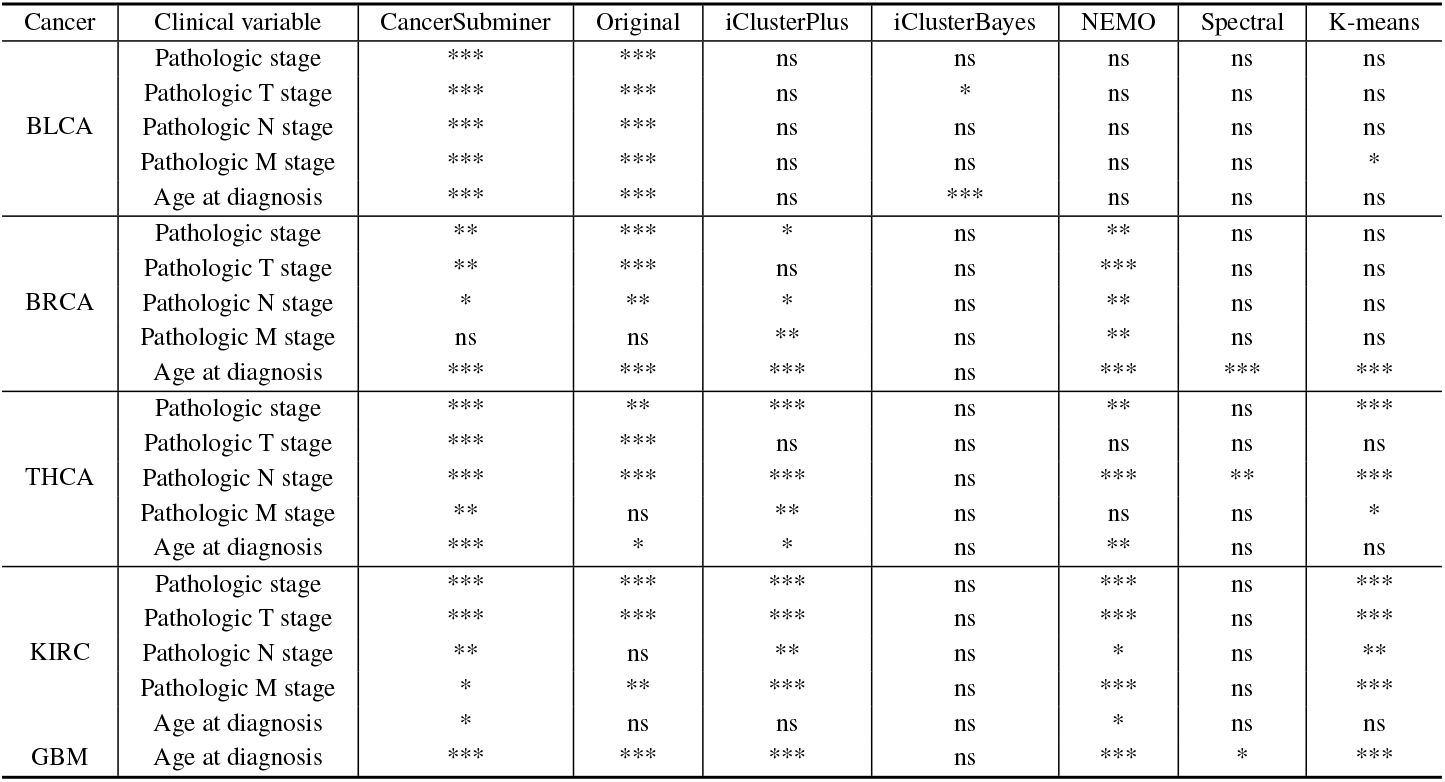
Clinical enrichment analysis results for identified subtypes, using statistical tests to assess between-subtype differences in clinical variables for each method.

For BLCA, our method identified significant associations for all clinical parameters, closely matching the performance of the established subtypes, whereas none of the comparison methods achieved similar results. In THCA and KIRC, CancerSubminer uncovered significant differences that were not observed with the established subtypes—specifically, pathologic M stage in THCA and pathologic N stage along with age at initial diagnosis in KIRC. These findings demonstrate that CancerSubminer not only preserves clinically relevant information captured during pre-training with established subtype annotations but also learns improved representations that refine and redefine subtypes, enabling clearer distinction of clinically meaningful groups.

### 4.4 Evaluating the role of reassignment in CancerSubminer

To evaluate the contribution of individual components in CancerSubminer, we conducted an ablation study on BRCA and THCA datasets. Specifically, we compared three configurations of the proposed framework: (i) without target-domain reassignment and without source-domain reassignment, (ii) without source reassignment, and (iii) the full model including all components. Performance was assessed using Silhouette score and DBI, with results summarized in Table x.

Across both cancer types, the full model consistently achieved the best performance, demonstrating the importance of incorporating both source and target reassignment strategies. For BRCA, the Silhouette score improved substantially compared to the PAM50 baseline, rising from 0.242 (TCGA-BRCA) and 0.180 (GSE72245) to 0.464 and 0.409, respectively, with the complete framework. Similarly, DBI values decreased markedly, indicating enhanced cluster compactness. For the additional BRCA target cohorts (GSE75067 and GSE69914), the full model also produced higher Silhouette scores and lower DBI compared to partial configurations, confirming its robustness across datasets.

For THCA, the proposed framework outperformed the original subtypes in both Silhouette score and DBI. The full model achieved the highest Silhouette score of 0.685 in TCGA-THCA, surpassing both the original subtypes (0.591) and the ablated variants. Comparable improvements were observed across external cohorts (e.g., Silhouette score of 0.637 in GSE97466 and 0.658 in GSE86961). While DBI varied depending on the dataset, the full model consistently delivered competitive or superior results relative to the ablated versions.

These results confirm that both source and target reassignment play complementary roles in refining subtype boundaries. Removing either component led to reduced clustering quality, whereas their joint inclusion provided the most consistent and robust performance across datasets. This demonstrates that CancerSubminer benefits from the synergy between pre-training with established subtypes and iterative refinement through reassignment strategies.

## 5 Discussions and Conclusion

This study introduces CancerSubminer, a methylation-based subtyping framework that integrates supervised learning, confidence-aware clustering, adversarial domain adaptation, and semi-supervised fine-tuning to address three persistent challenges in cancer subtyping: reliance on reference labels that overlook features outside the training distribution, cohort-specific batch effects, and the trade-off between maintaining known subtype integrity and uncovering new structure. Across five cancer types, CancerSubminer consistently improved cluster compactness and separation relative to established annotations and leading baselines. The derived subtypes achieved significant survival stratification and stronger associations with clinical parameters, demonstrating that the framework captures clinically meaningful heterogeneity beyond existing definitions.

A key strength of CancerSubminer lies in its ability to refine canonical labels while remaining receptive to patterns previously underrepresented in existing classifications. For example, in thyroid carcinoma, conventional subtypes showed less association with patient outcomes, whereas CancerSubminer revealed subgroups with significant survival differences, indicating that methylation-derived signals preserve prognostic relevance overlooked by earlier subtyping systems. Similar improvements were observed in breast and bladder cancers, where the framework produced more coherent clusters and stronger clinical associations than competing approaches. These results show that the sequential process of supervised pretraining, confidence-based reassignment, and domain alignment effectively extracts structured signal aligned with clinical outcomes.

Outcome-level validation through Kaplan–Meier analysis further supports this conclusion. Subtypes derived by CancerSubminer achieved statistically significant survival separation in all five cancer types (log-rank p < 0.05), whereas other methods produced inconsistent results. The persistence of prognostic distinctions in a methylation-defined subtype space suggests that the learned representations encode information with independent predictive value, reinforcing their potential clinical utility.

The ablation study clarifies the contribution of each module. Confidence-based reassignment on both the source and target domains proved crucial for stabilizing subtype boundaries. Removing either component degraded cluster quality, while their combination yielded the most consistent and clinically interpretable outcomes. These findings indicate that ambiguous samples near decision margins introduce noise that can be reduced when reassignment is guided by high-confidence peers, while target-aware reassignment prevents error propagation from uncertain pseudo-labels.

Equally important, adversarial domain adaptation and subtype alignment effectively minimized technical variation without diluting subtype signal. These components promoted domain-invariant embeddings in which subtype clusters spanned across cohorts rather than fragmenting within them—a prerequisite for reproducible subtype discovery and cross-institutional translation. Visualization further supports this interpretation, showing reduced batch-driven separation while maintaining clear subtype distinctions.

Although multi-omics integration can increase discovery power, clinical implementation often faces constraints due to missing modalities and cost. By demonstrating strong performance using methylation data alone, CancerSubminer provides a practical and scalable approach for institutions with established methylation workflows. The modular design also enables future expansion to multi-omics inputs while preserving its core reassignment, alignment, and clustering strategies, supporting flexible adaptation as additional data types become available.

Several limitations suggest directions for future work. External validation with prospectively collected, diverse cohorts will be necessary to confirm generalizability and mitigate spectrum bias. Sensitivity analysis of sample sizes would help characterize stability across different data regimes. In addition, while CancerSubminer identifies potential novel subtypes, experimental and functional validation—such as pathway enrichment or integration with transcriptomic and proteomic data—will be required to substantiate their distinctiveness.

In summary, CancerSubminer advances methylation-based subtyping by jointly resolving label bias, batch effects, and the discovery–preservation tension that often limits translational application. It consistently produces compact, well-separated subtypes with improved prognostic stratification and stronger clinical associations across diverse cancer types. These capabilities position CancerSubminer as a practical and extensible foundation for precision oncology workflows that rely on single-omics data, heterogeneous cohorts, and the need for reproducible subtyping results.

