## Supplementary Material for "CancerSubminer: an integrated framework for cancer subtyping using supervised and unsupervised learning on DNA methylation profiles"

**Supplementary Material S1**. Silhouette scores for K-means and Spectral clustering across different numbers of clusters. Bold values indicate the highest score used to determine the optimal number of clusters.

| | | | | | | |
|---|---|---|---|---|---|---|
| BLCA | # of clusters | 3 | 4 | 5 | 6 | 7 |
| | K-means | 0.3853 | **0.3882** | 0.3107 | 0.3078 | 0.1981 |
| | Spectral | 0.0323 | **0.0782** | -0.1654 | -0.0426 | -0.2305 |
| BRCA | # of clusters | 3 | 4 | 5 | 6 | 7 |
| | K-means | **0.2756** | 0.2681 | 0.2457 | 0.2142 | 0.1807 |
| | Spectral | -0.1716 | **-0.0067** | -0.0825 | -0.0772 | -0.3486 |
| GBM | # of clusters | 3 | 4 | 5 | 6 | 7 |
| | K-means | 0.1942 | **0.3917** | 0.3458 | 0.2984 | 0.3020 |
| | Spectral | **-0.0133** | -0.2196 | -0.1480 | -0.0698 | -0.4893 |
| KIRC | # of clusters | 2 | 3 | 4 | 5 | 6 |
| | K-means | 0.0899 | **0.5583** | 0.5103 | 0.3887 | 0.2844 |
| | Spectral | 0.0155 | -0.0224 | **0.0868** | -0.2585 | -0.4535 |
| THCA | # of clusters | 3 | 4 | 5 | 6 | 7 |
| | K-means | **0.3687** | 0.2601 | 0.2366 | 0.1667 | 0.2018 |
| | Spectral | **0.0002** | -0.0571 | -0.0615 | -0.2168 | -0.0116 |

**Supplementary Material S2**. UMAP visualization of source and target datasets for BRCA, GBM and KIRC. (a) UMAP plot of the uncorrected dataset, colored by batch. (b) UMAP plots of features extracted by CancerSubminer, colored by batch (left) and by identified cancer subtypes (right). (c) UMAP plots of subtype assignments from comparison methods, including NEMO, iClusterPlus, iClusterBayes, Spectral Clustering, K-means, and the original dataset annotations.

#### BRCA

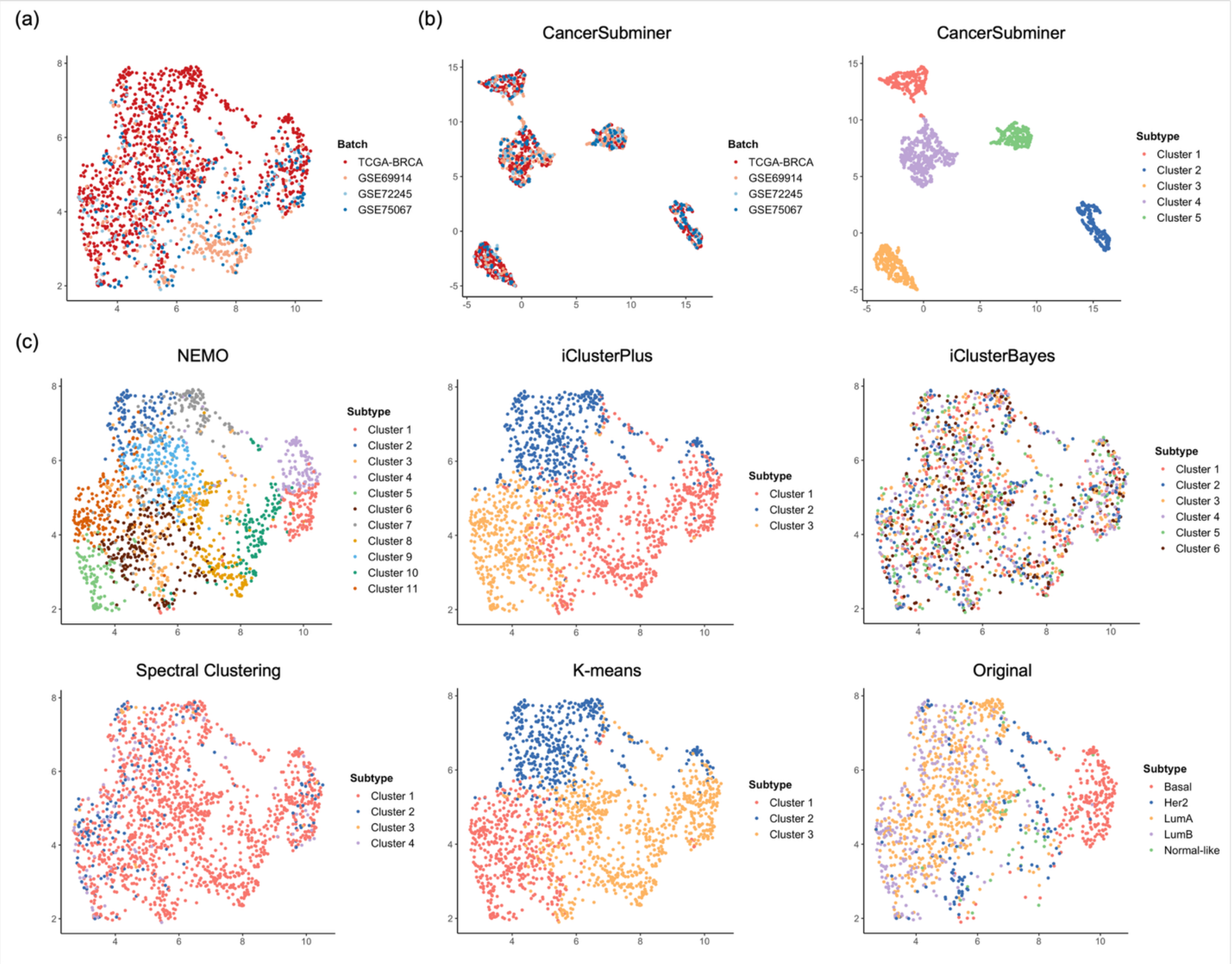

### GBM

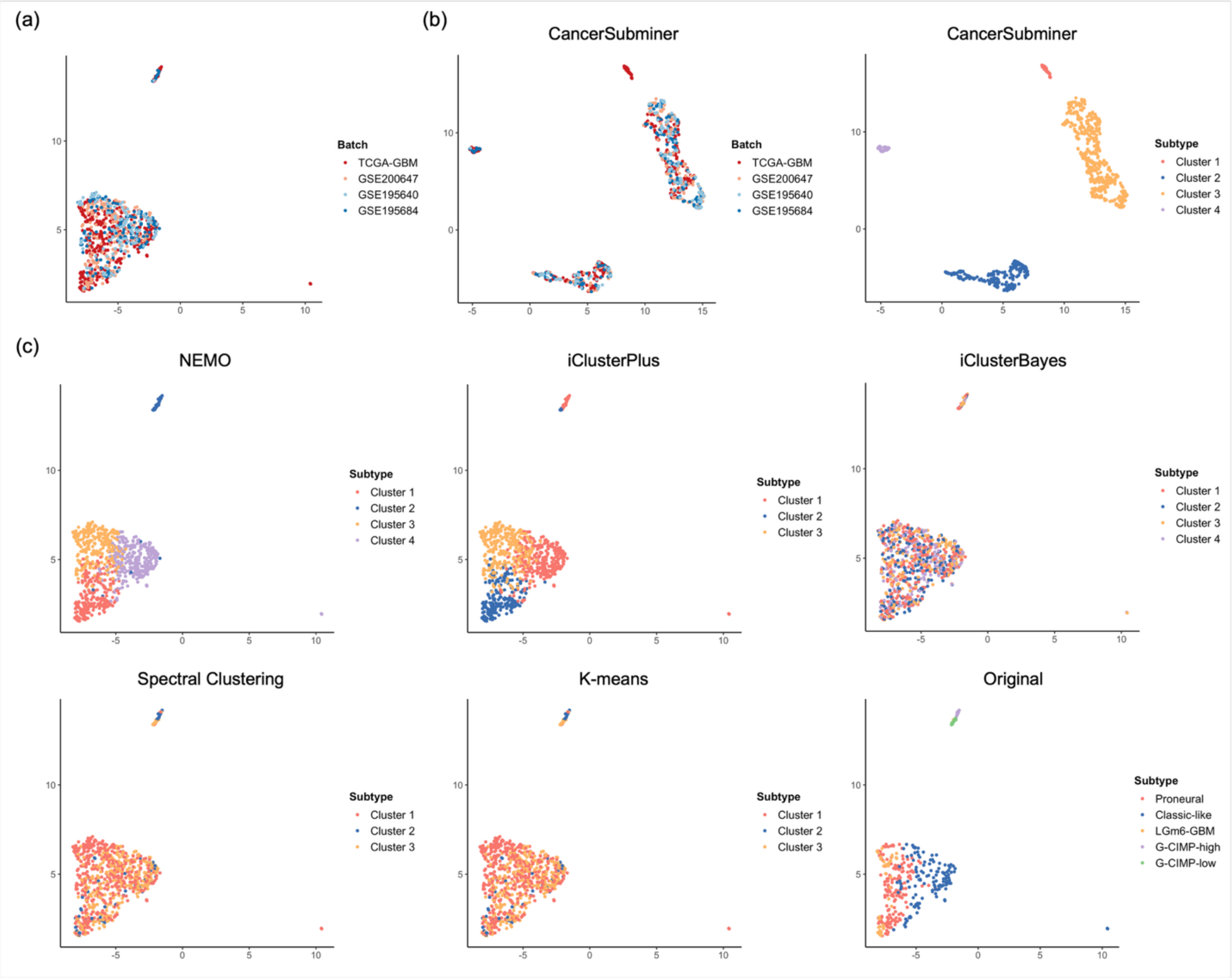
(a)
Batch
TCGA-GBM
GSE200647
GSE195640
GSE195684
(b)
CancerSubminer
Batch
TCGA-GBM
GSE200647
GSE195640
GSE195684
CancerSubminer
Subtype
Cluster 1
Cluster 2
Cluster 3
Cluster 4
(c)
NEMO
Subtype
Cluster 1
Cluster 2
Cluster 3
Cluster 4
iClusterPlus
Subtype
Cluster 1
Cluster 2
Cluster 3
iClusterBayes
Subtype
Cluster 1
Cluster 2
Cluster 3
Cluster 4
Spectral Clustering
Subtype
Cluster 1
Cluster 2
Cluster 3
K-means
Subtype
Cluster 1
Cluster 2
Cluster 3
Original
Subtype
Proneural
Classic-like
LGm6-GBM
G-CIMP-high
G-CIMP-low

### KIRC

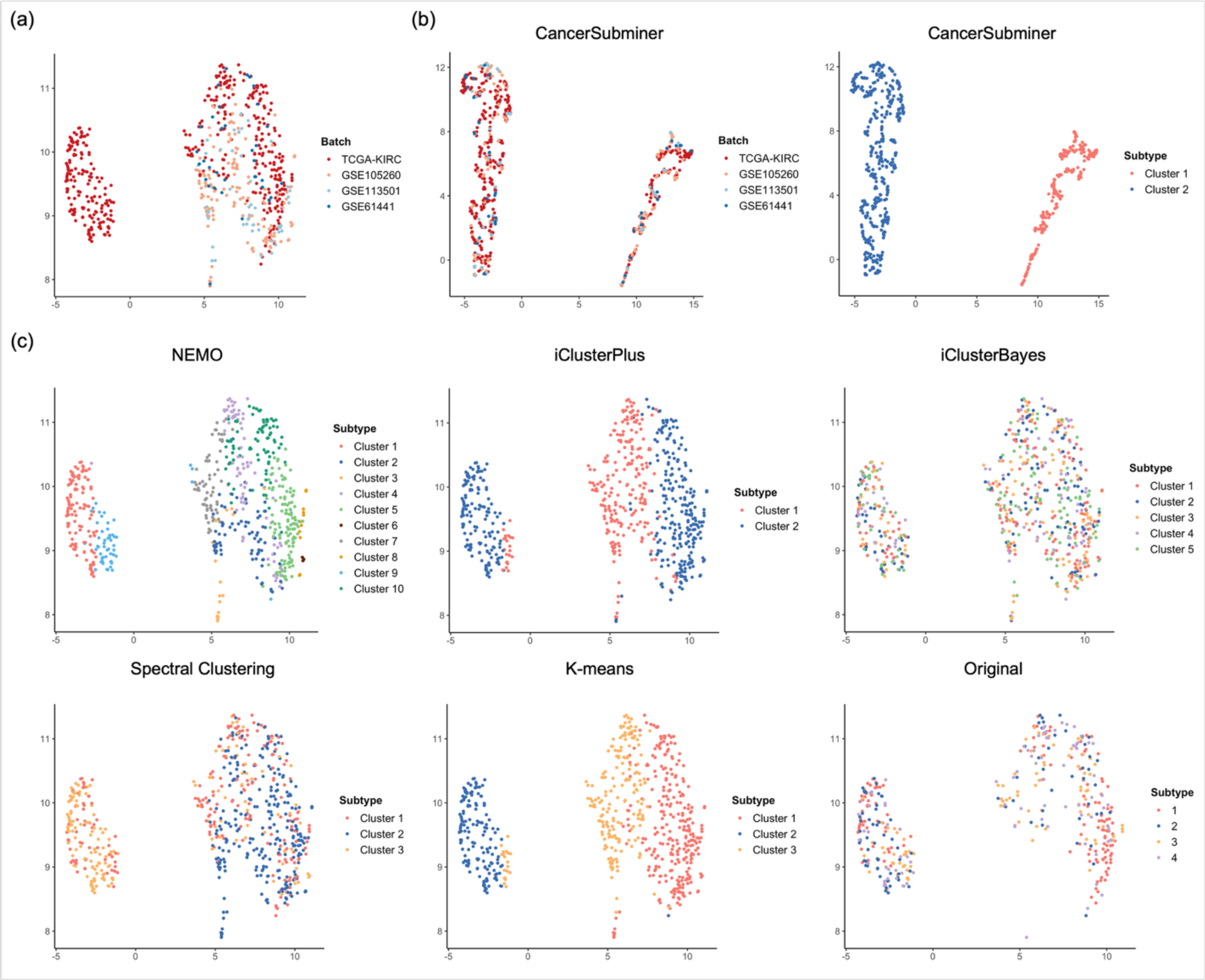
(a)
Batch
TCGA-KIRC
GSE105260
GSE113501
GSE61441
(b)
CancerSubminer
Batch
TCGA-KIRC
GSE105260
GSE113501
GSE61441
CancerSubminer
Subtype
Cluster 1
Cluster 2
(c)
NEMO
Subtype
Cluster 1
Cluster 2
Cluster 3
Cluster 4
Cluster 5
Cluster 6
Cluster 7
Cluster 8
Cluster 9
Cluster 10
iClusterPlus
Subtype
Cluster 1
Cluster 2
iClusterBayes
Subtype
Cluster 1
Cluster 2
Cluster 3
Cluster 4
Cluster 5
Spectral Clustering
Subtype
Cluster 1
Cluster 2
Cluster 3
K-means
Subtype
Cluster 1
Cluster 2
Cluster 3
Original
Subtype
1
2
3
4

**Supplementary Material S3**. Kaplan–Meier survival analysis of BRCA subtypes identified from the GSE75067 dataset, with log-rank test results comparing CancerSubminer against baseline clustering methods.

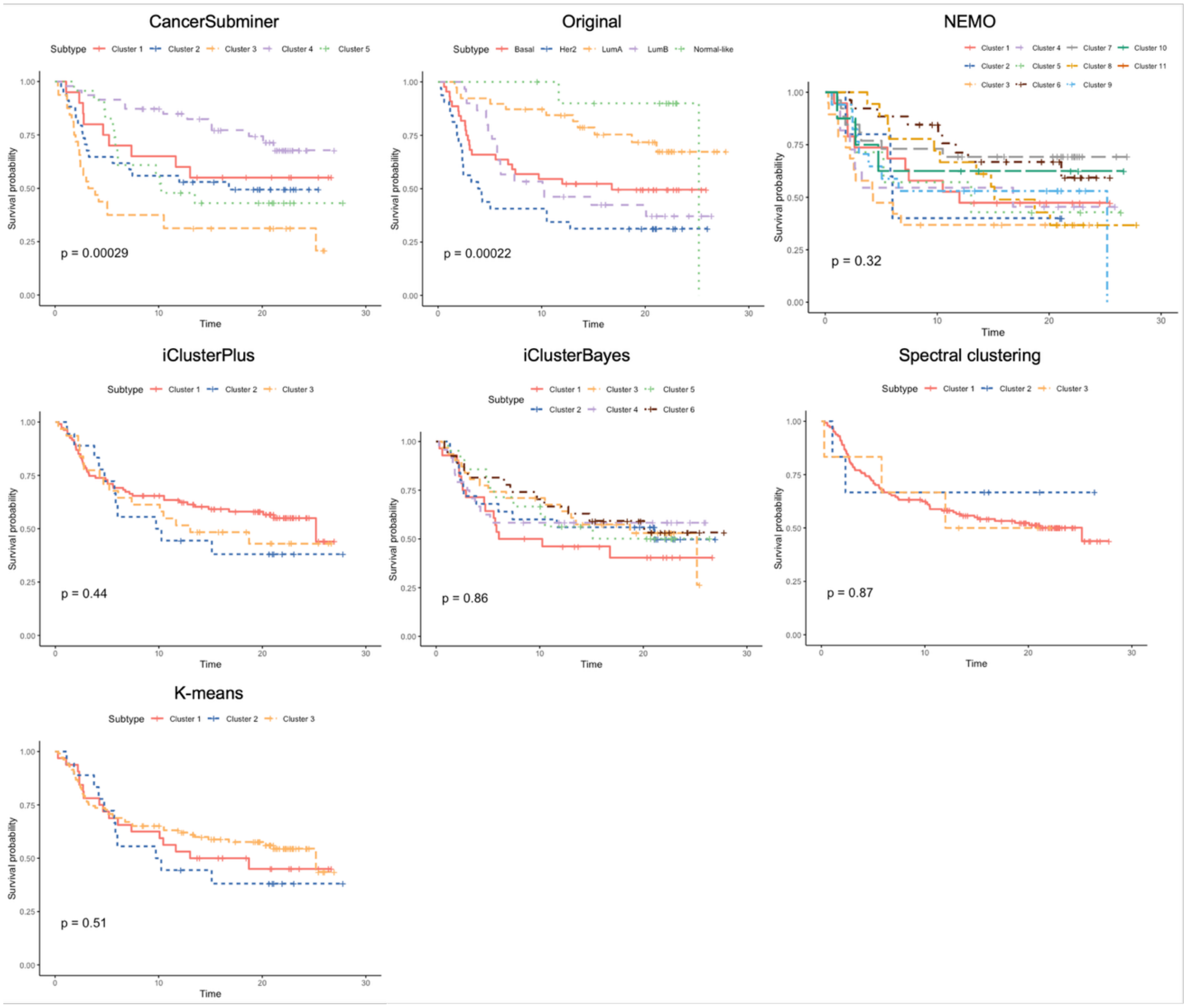

**Supplementary Material S4**. Kaplan–Meier survival analysis of BLCA, GBM, and KIRC subtypes identified from the TCGA dataset, with log-rank test results comparing CancerSubminer against baseline clustering methods.

### BLCA

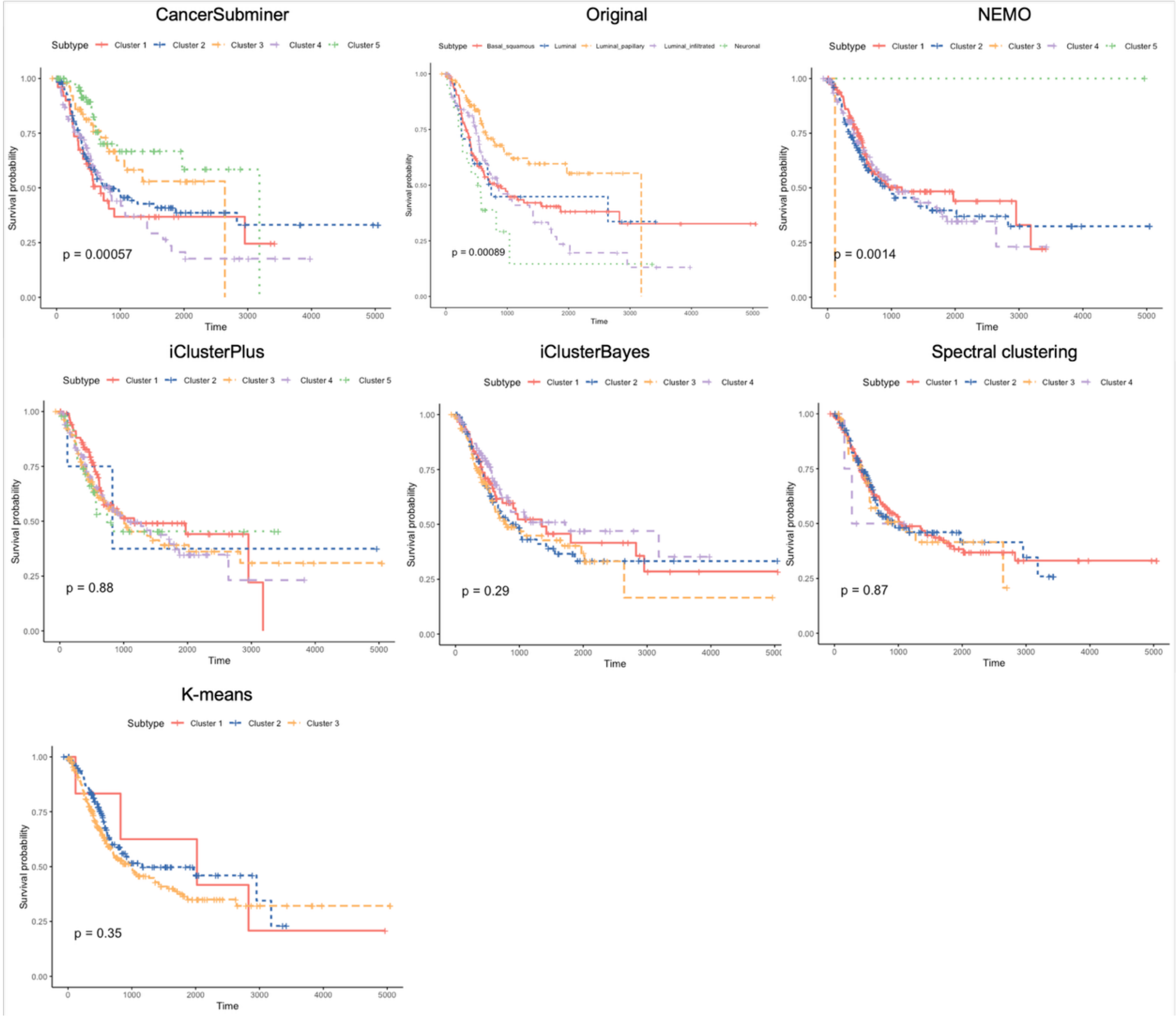

### GBM

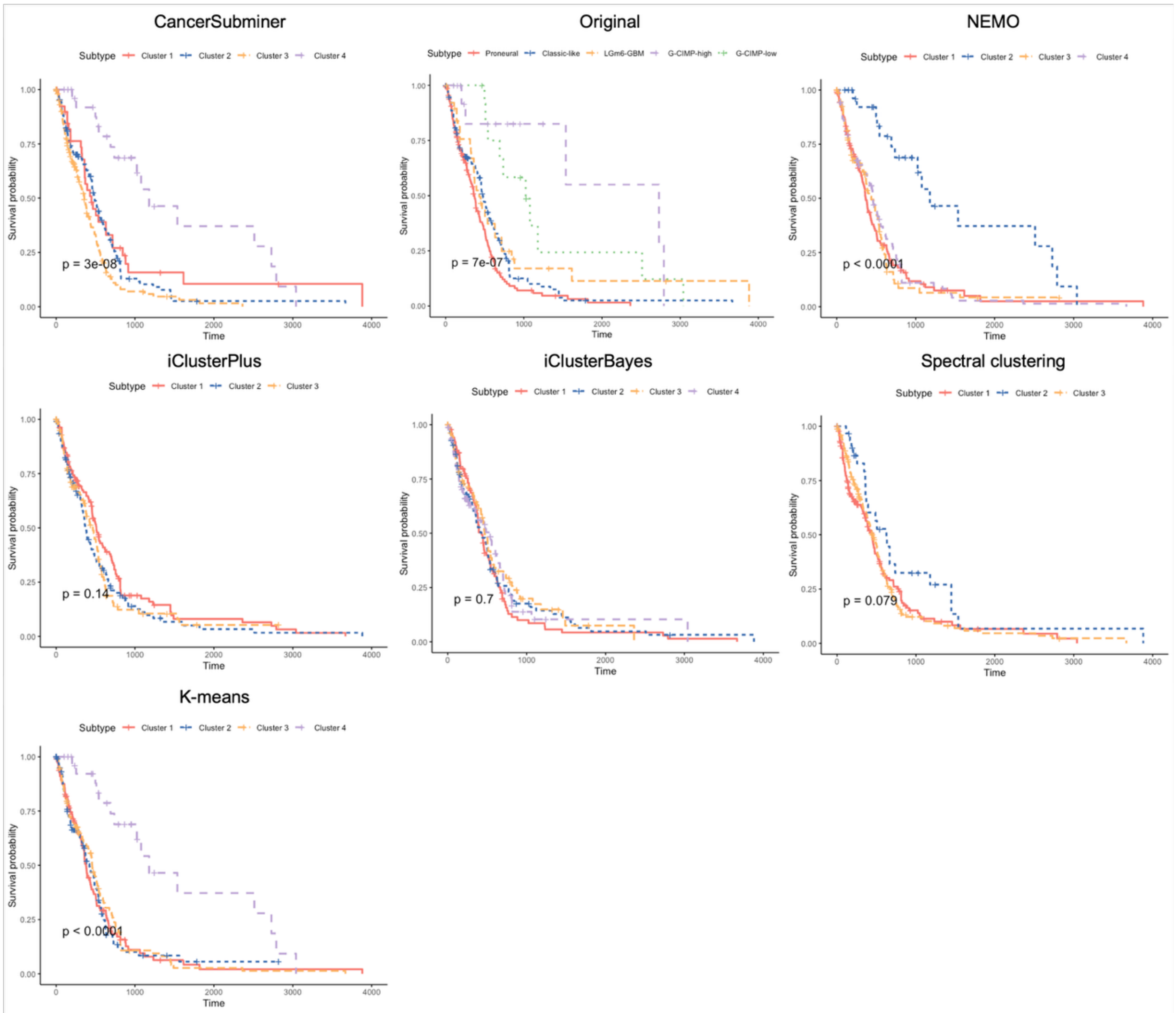

CancerSubminer
Subtype Cluster 1 Cluster 2 Cluster 3 Cluster 4
p = 3e-08
Original
Subtype Proneural Classic-like LGm6-GBM G-CIMP-high G-CIMP-low
p = 7e-07
NEMO
Subtype Cluster 1 Cluster 2 Cluster 3 Cluster 4
p < 0.0001
iClusterPlus
Subtype Cluster 1 Cluster 2 Cluster 3
p = 0.14
iClusterBayes
Subtype Cluster 1 Cluster 2 Cluster 3 Cluster 4
p = 0.7
Spectral clustering
Subtype Cluster 1 Cluster 2 Cluster 3
p = 0.079
K-means
Subtype Cluster 1 Cluster 2 Cluster 3 Cluster 4
p < 0.0001
Survival probability
Time

### KIRC

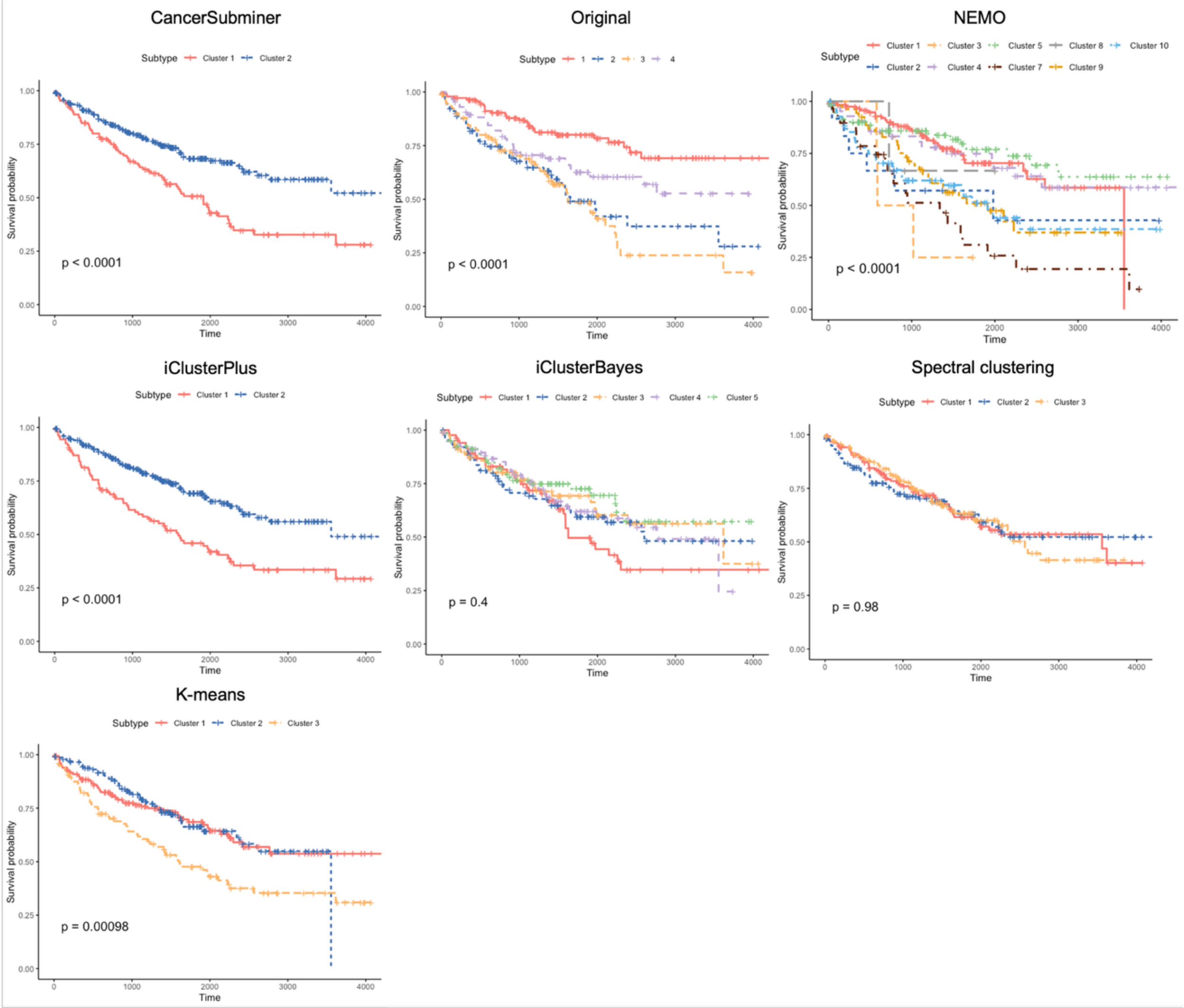

CancerSubminer
Subtype Cluster 1 Cluster 2
p < 0.0001
Original
Subtype 1 2 3 4
p < 0.0001
NEMO
Subtype Cluster 1 Cluster 3 Cluster 5 Cluster 8 Cluster 10 Cluster 2 Cluster 4 Cluster 7 Cluster 9
p < 0.0001
iClusterPlus
Subtype Cluster 1 Cluster 2
p < 0.0001
iClusterBayes
Subtype Cluster 1 Cluster 2 Cluster 3 Cluster 4 Cluster 5
p = 0.4
Spectral clustering
Subtype Cluster 1 Cluster 2 Cluster 3
p = 0.98
K-means
Subtype Cluster 1 Cluster 2 Cluster 3
p = 0.00098
Survival probability
Time
